# Active transport of brilliant blue FCF across the *Drosophila* midgut and Malpighian tubule epithelia

**DOI:** 10.1101/771675

**Authors:** Dawson B.H. Livingston, Hirva Patel, Andrew Donini, Heath A. MacMillan

**Affiliations:** Department of Biology, Carleton University, Ottawa, Canada, K1S 5B6; Department of Biology, York University, Toronto, Canada, M3J 1P3

**Keywords:** Organic anion, paracellular barriers, barrier dysfunction, renal function, cold tolerance, FD&C blue dye no. 1

## Abstract

Under conditions of stress, many animals suffer from epithelial barrier disruption that can cause molecules to leak down their concentration gradients, potentially causing a loss of organismal homeostasis, further injury or death. *Drosophila* is a common insect model, used to study barrier disruption related to aging, traumatic injury, or environmental stress. Net leak of a non-toxic dye (Brilliant blue FCF) from the gut lumen to the hemolymph is often used to identify barrier failure under these conditions, but *Drosophila* are capable of actively transporting structurally-similar compounds. Here, we examined whether cold stress (like other stresses) causes Brilliant blue FCF (BB-FCF) to appear in the hemolymph of flies fed the dye, and if so whether *Drosophila* are capable of clearing this dye from their body following chilling. Using *in situ* midgut leak and transport assays as well as Ramsay assays of Malpighian tubule transport, we tested whether these ionoregulatory epithelia can actively transport BB-FCF. In doing so, we found that the *Drosophila* midgut and Malpighian tubules can mobilize BB-FCF via an active transcellular pathway, suggesting that elevated concentrations of the dye in the hemolymph may occur from increased paracellular permeability, reduced transcellular clearance, or both.

**Summary Statement:** *Drosophila* are able to actively secrete Brilliant blue FCF, a commonly used marker of barrier dysfunction

## Introduction

Epithelial barriers are an important constituent of all metazoan life on earth and are critical in the maintenance of homeostasis. In the gut, epithelial barriers play a major role in regulating rates of nutrient absorption; ion and water secretion and absorption; the clearance of wastes (Salvo Romero et al., 2015); and protect organisms from potential threats such as infection and dehydration (Presland and Jurevic, 2002).

*Drosophila* are a common model of digestive and renal physiology (Dow & Romero, 2010; Buchon et al., 2013; Miguel-Aliaga et al., 2018). The gut of *Drosophila* is divided into three distinct regions; the foregut, midgut and hindgut. The foregut primarily functions in food storage and regulates the passage of food into the midgut. The insect midgut is largely analogous to the small intestine in humans, and is the primary site of food digestion, and nutrient absorption (Buchon et al., 2013). From the midgut, digested material enters the hindgut, which functions in water resorption prior to defecation. The Malpighian tubules are blind-ended tubes that branch from the midgut-hindgut junction, and are a single cell layer thick (Demerec, 1950) The Malpighian tubules are responsible for the production of primary urine containing water, ions, toxins, and waste products (Dow, 2009; Buchon et al., 2013). *Drosophila* have four Malpighian tubules; two anterior and two posterior, which all function in ion and water transport (Denholm and Skaer, 2009; Chintapalli et al., 2012).

In addition to its critical role in facilitating our understanding of gut and renal physiology, *Drosophila* are also used to probe the roles of epithelial barriers in aging and disease (Dow et al., 1994; Rera et al., 2012; Rodan et al., 2012; He & Jasper, 2014; Clark et al., 2015; Dambroise et al., 2016). Advanced age, infections and thermal stress, for example, are all known to cause a loss or decrease in epithelial barrier function (Jasper, 2015; MacMillan et al., 2017), and this barrier dysfunction is thought to contribute to ageing phenotypes and/or stress-induced injury or death (Katzenberger et al., 2015; Cerqueira César Machado and Pinheiro da Silva, 2016). In most of these studies, barrier dysfunction in fruit flies is identified and quantified using the “Smurf assay” (Rera et al., 2011, Martins et al., 2018). In this assay, flies are fed a food containing a high concentration of a non-toxic organic dye called Brilliant blue FCF (BB-FCF). Flies that are healthy appear to retain the dye in their gut lumen, but those that have some form of intestinal epithelial dysfunction turn blue (Rera et al., 2011). It is presumed that the presence of these Smurf phenotypes is a direct result of BB-FCF leaking down its concentration gradient (i.e. passively diffusing) from the gut lumen into the hemolymph because of barrier disruption, which will eventually make the fly appear blue (Rera et al., 2011).

When exposed to cold temperatures, *D. melanogaster* become less capable at regulating hemolymph osmolarity and ion balance, and as a result, hyperkalemia causes cell depolarization and cell death (MacMillan et al., 2015a). This inability to maintain ion and water balance in the cold has been attributed, in part, to a susceptibility of the renal system to chilling (Overgaard and MacMillan, 2017). Specifically, low temperatures suppress ionomotive enzyme activity in the Malpighian tubules and gut epithelia, such that a net leak of Na^+^ and water from the hemolymph to the gut causes a toxic increase in concentration of K^+^ in the hemolymph (Overgaard and MacMillan, 2017), This cascade of issues can be largely prevented in *Drosophila* given a period of cold acclimation (MacMillan et al., 2015b; Yerushalmi et al., 2018). When ion balance is disturbed, recovery of K^+^ balance (and thus cell membrane potential) is thought to be particularly important to the recovery of neuromuscular function following rewarming (MacMillan et al., 2012; Andersen and Overgaard, 2019). The ability of insects to recover can be quantified by measuring the time required for an insect to stand following chilling (termed chill coma recovery time; CCRT) (Macdonald et al., 2004; Findsen et al., 2014). Like other stressful conditions, cold exposures also cause gut barrier dysfunction (MacMillan et al., 2017), but this cold-induced barrier failure was described using another common paracellular probe (dextran) and has previously not been investigated using BB-FCF.

Some solutes are able to pass through the epithelial cells of an insect gut, while others can’t (Huang et al., 2015). Large polar organic molecules (like BB-FCF or dextran) cannot easily pass through cell membranes, but there are two primary routes by which they could potentially cross epithelial barriers; transcellular and paracellular transport. Transcellular transport occurs across cell membranes via active (i.e. energy-consuming) or passive routes (O’Donnell & Maddrell, 1983). Many complex organic compounds have been shown to be actively (e.g. para-aminohippuric acid, tetraethylammonium) (Linton & O’Donnell, 2000; Rheault & O’Donnell, 2004), or passively (e.g. urea, sucrose) (Maddrell & Gardiner, 1974) transported across *Drosophila* epithelia. Many inorganic ions and compounds are actively transported at high rates by the gut and renal epithelia of insects (e.g. phosphate, Mg^2+^) (Maddrell & O’Donnell, 1992). Passive diffusion of a molecule down its concentration gradient can occur in the transcellular manner (e.g. through suitable channels) provided the molecule has a low molecular weight and is soluble in the hemolymph. By contrast, primary active transport occurs transcellularly through dedicated transporters and secondary active transport can mobilize molecules up or down a concentration gradient by coupling concentration gradients through exchangers (Maddrell & O’Donnell, 1992; Rheault & O’Donnell, 2004).

Paracellular transport is defined as the transport of molecules/solutes across the epithelial cell layer, specifically between the cells via the intercellular spaces (O’Donnell & Maddrell, 1983). In insects, paracellular permeability is regulated by the presence of septate junctions (SJ), protein complexes analogous to vertebrate tight junctions that play an important role in selectivity of passage (O’Donnell & Maddrell, 1983; Jonusaite et al., 2017). SJs act as permeability barriers to the free diffusion of molecules, and thereby control the rate at which molecules can leak down their concentration gradient from one side of an epithelium to the other (Banerjee et al., 2006; Ganot et al., 2014; Jonusaite et al., 2016). There are two types of these junctions in insects, pleated and smooth SJs; while pleated SJs are typically found in the salivary glands, epidermis and the photoreceptors of the flies, smooth SJs are primarily found in endodermal tissues like the midgut and Malpighian tubules (Hall and Ward, 2016; Jonusaite et al., 2017).

The mode of transport of BB-FCF through the *Drosophila* gut epithelia is unknown, and despite its use in understanding epithelial health and disease, whether or not flies can actively transport this dye remains untested. Importantly, other organic compounds can be actively transported by the *Drosophila* midgut and Malpighian tubules, including TEA, Texas red and methotrexate (Rheault & O’Donnell, 2004; O’Donnell & Leader, 2006; O’Donnell, 2009). Texas red (625.12 g mol^-1^), for example, is of a similar size to brilliant blue (792.85 g mol^-1^), and is transported across the Malpighian tubule epithelia via the action of a sodium-independent active transport mechanism (Leader and O’Donnell, 2005). This active transport has previously been confirmed by experimentally saturating the putative transporter with high concentrations of Texas red (Leader and O’Donnell, 2005).

In the present study, we hypothesized that 1) Like other stressors, cold stress can cause the Smurf phenotype in *Drosophila*, but that this outcome could be due to either barrier failure or a reduced capacity for active transport in the cold because 2) the gut and Malpighian tubule epithelia of *Drosophila* are innately capable of actively secreting BB-FCF. We used dye leak and clearance assays using whole animals and isolated organs to test whether and how flies transport the dye and found that cold stress does indeed cause the Smurf phenotype in *Drosophila* and that Smurf flies are somewhat more likely to die from cold stress. We argue that Smurf assays, however, cannot distinguish between a decrease of dye transport rates and a failure to maintain epithelial barriers under conditions of stress, as the midgut and tubules are continuously clearing the dye from the hemolymph via active transcellular transport.

## Materials and Methods

### Animal Husbandry

The line of *Drosophila melanogaster* used in this study were derived from isofemale lines established from collections from London, Ontario and Niagara on the Lake, Ontario in 2007 (Marshall and Sinclair, 2009). *D. melanogaster* were reared in 200 mL bottles that contained ∼50 mL of Bloomington *Drosophila* medium (a cornmeal, corn syrup, agar and yeast-based diet). FD&C Blue dye #1 (brilliant blue FCF, BB-FCF) was a generous gift from Calico Food Ingredients Ltd. (Kingston, ON). The flies used in our cold tolerance assays were kept at 25°C with a 14 hr: 10 hr light:dark cycle in a MIR-154-PA incubator (Panasonic Corporation, Kadoma, Japan). Adult flies were transferred to a fresh bottle with food to lay eggs for two hours to ensure a consistent number (∼200) of progeny per bottle. Following adult emergence (approximately ten days after the eggs were laid), the flies were collected in 40 mL vials containing 7 mL of food medium. The flies were sexed on a CO_2_ pad three days post-adult-emergence (day 13 of their life cycle), this was done in less than 5 min to reduce the time flies are exposed to the CO_2_. All of the experiments were completed on adult females, seven days after final ecdysis.

### Survival Assay

A total of 250 flies were used to quantify survival rates following cold stress. Dyed medium was prepared using the standard rearing medium, but with BB-FCF added at a concentration of 2.5% (w/v) (Rera et al., 2012). Flies were maintained on dyed medium for nine hours and then transferred into empty 40 mL vials. Control flies (n=50; ∼10 per vial) were observed at this point for the Smurf phenotype. A fly was counted as Smurf when dye coloration was observed outside of the digestive tract (observed using a microscope, without dissection), while they lay in ventral position. After observation the control flies were transferred to new vials containing regular Bloomington food for 24 h before survival was assessed. The rest of the vials were put in a Styrofoam container with a mixture of water and ice (0°C). After 6, 12, 18 and 24h at 0°C, subsamples of 50 flies were removed from the cold stress. After observation for the Smurf/non-Smurf phenotypes, the flies were transferred to new vials containing standard Bloomington food while separating the Smurf flies from the Non-Smurf flies into groups of ∼10 flies/vial. The vials with the recovering flies were left on their side for 24h before survival was assessed. Flies then able to stand and walk were counted as alive, and flies not moving or moving but unable to stand and walk counted as dead.

### Chill coma recovery time (CCRT)

Flies were maintained on dyed medium (prepared as described above) for 9 h before individual flies were transferred into empty 4 mL glass screw-top vials. The vials were put into a plastic bag (10 vials/bag) and submerged in an ice-water mixture (0°C). After 12 h, all of the vials were removed from the ice-water bath. Each fly was immediately assessed to determine whether or not they had a Smurf phenotype. All of the vials were then arranged on a sheet of paper on a laboratory bench at room temperature (23°C). All of the flies were observed for 1 h to record CCRT. The CCRT was recorded as the time it took for the fly to stand up on all six legs, and the flies that did not stand within the hour were considered to have suffered chilling injury.

### Cold acclimation

Thermal acclimation over a period of days can strongly influence cold tolerance in *D. melanogaster*, and cold acclimated flies better maintain barrier function in the cold (MacMillan et al., 2017). To test whether cold acclimation prevented flies from having a Smurf phenotype, we acclimated flies to one of two temperatures and examined their ability to retain the dye in their gut. Flies (reared as described above) were collected on the day of adult emergence and sorted by sex under short CO_2_ anaesthesia (< 5 minutes) before being transferred to either 10°C or 25°C. At six days of age (post adult ecdysis), warm- and cold-acclimated flies were transferred to fresh vials of their standard medium containing 2.5% (w/v) of BB-FCF. The flies were separated (without anaesthesia) into groups of 20-26 flies per vial (four vials for each acclimation group) and the vials were placed in an ice-water mixture (at 0°C) for 16 h. After removal, and while the flies were still in chill coma, they were placed under the view of a dissecting microscope and the number of Smurf flies and non-Smurf flies were counted.

### Dye clearance assay

Dyed food medium was prepared as described above, and flies were maintained on the medium for 9 h after which they were then transferred to empty 40 mL vials. Control flies were observed at this point and scored for the Smurf phenotype. After observation, these control flies were transferred to 40 mL vials with 7 mL of regular Bloomington media for 12 hours before the ability of the flies to clear the dye from the gut could be seen. The remainder of the flies in the empty vials were placed in an ice bath at 0°C for 12 h. After examination of the flies for Smurf phenotype (observed using a microscope without dissection), the Smurfs were then placed into 40 mL vials with 7 mL of standard Bloomington media for 30 h, after which they were re-scored for the Smurf phenotype to assess their ability to clear the dye from the hemolymph. Photos of representative Smurf flies were taken directly following the cold stress using an Axiocam 105 color microscope camera (Zeiss, Oberkochen, Germany). Photos of the same flies were then taken for direct phenotypic comparison.

### Midgut dye leak and transport assay

To quantify the effects of cold exposure on the leak of BB-FCF across the *Drosophila* midgut, and whether the midgut is capable of transporting this dye, an *in situ* barrier function/transport assay was developed. Flies that had been fed on diet containing BB-FCF (as above) were dissected under paraffin oil such that the gut was surrounded by the oil but remained attached to the anterior (head and thorax) and posterior (abdomen) ends of the animal (both of which were held in place using minute dissecting pins). A single 2 µL droplet of *Drosophila* saline (117.5 mM NaCl; 20 mM KCl; 2 mM CaCl_2_; 8.5 mM MgCl_2_; 10.2 mM NaHCO_3_; 4.3 mM NaH_2_PO_4_; 15 mM MOPS; 20 mM glucose; 10 mM glutamine; pH 7.0) was placed on the posterior midgut. The preparation was then either left at 23°C, or was transferred to 0°C by placing it on the surface of an ice water mixture within an insulated container. After 3h, a 1 µL sample of the droplet on the gut was taken and mixed with 100 µL of milliQ water in a 96-well microplate. Absorbance was read at 630 nm and concentrations of BB-FCF that had leaked into the droplet from the gut were quantified by reference to an appropriately ranged BB-FCF standard curve. These values were then expressed relative to the length of the midgut inside the droplet in each preparation. To determine whether the *Drosophila* midgut could actively transport BB-FCF up a concentration gradient, we conducted the same assay at 23°C, but with a 2 µL droplet of saline containing a known concentration of the dye (75 µM), well below that of the diet.

### Ramsay Assays

To test whether the Malpighian tubules can transport BB-FCF, and whether this transport is likely to be carrier-mediated, we used a modified Ramsay assay procedure, similar to that described by Rheault and O’Donnell (Rheault and O’Donnell, 2004). The flies used for the Ramsay assays were reared under the same conditions with the exception that the light:dark cycle was 12 hr:12 hr (simply because of available rearing space). Individual flies were briefly aspirated into ethanol and the exoskeleton was carefully dissected away to reveal the gut and Malpighian tubules. The Malpighian tubules were excised by cutting at the base of the ureter using a 3G needle. Using a thin pulled glass rod, the pair of tubules was transferred to a saline droplet in a dish containing heavy paraffin oil. The control droplets submerged in the oil were 10 µL of 1:1 mixture of *Drosophila* saline (described above) and Schneider’s Insect Medium (Sigma-Aldrich S0146, St. Louis, USA). The experimental droplets contained the same 1:1 mixture mentioned above, but had BB-FCF added to obtain a range of dye concentrations between 0.0126 µM and 25.2 µM (n=5-15 tubules per concentration). One tubule was wrapped around a pin outside of the droplet, such that the ureter was half-way between the droplet and the pin, and the other tubule remained in the droplet. The droplet that formed at the ureter was removed after 15 min, and droplet volumes were approximated using droplet diameter, which was measured using an ocular micrometer (assuming a sphere when the droplet was suspended in the oil). The droplets were then collected using a 2 µL micropipette and transferred to an empty microcentrifuge tube. A 3 µL volume of *Drosophila* saline was added to each sample in order to separate the aqueous fluid from any oil collected along with the droplet. After brief centrifugation, a 2 µL aliquot of aqueous layer was transferred to a Take3 microwell plate (BioTek, Winooski, USA) and the absorbance was read using the Biotek Cytation 5 Imaging Reader at 630 nm. The absorbance reading was then compared to a standard curve to determine the concentration of BB-FCF, which was then used to calculate the original concentration in the primary urine.

### Data Analysis

All data analysis was completed using R (version 3.5.2). Linear regression was used to assess whether the tendency for flies to adopt a Smurf phenotype increased with cold exposure duration, and survival following cold stress was analyzed using a generalized linear model with a binomial error distribution (with phenotype and time included as factors). A Welch’s two sample t-test were used to determine whether there was a difference in CCRT between Smurfs and non-Smurfs, and whether cold acclimation altered the tendency of flies to become Smurfs. The effect of temperature on the leak of BB-FCF from the midgut was examined using a one-way ANOVA. To analyze our Ramsay assay data, we fit a Michaelis-Menton type model to the relationship between the concentration of BB-FCF bathing the Malpighian tubules and the rate of dye flux (nmol min^-1^) through the tubules using the nls() function. This model was used to determine the maximal rate of dye clearance (J_max_) and the saline concentration at which dye transport rates were 1/2 the maximal rate (K_t_).

## Results

### Low temperature survival and chill coma recovery

Increased cold exposure times increased the likelihood of a fly becoming a Smurf, and decreased the probability of survival following chilling. We found a significant linear relationship between the duration of exposure to 0°C and the proportion of the flies that turned blue (i.e. became Smurfs; *P* = 0.049; Fig. 1A). Longer exposures to 0°C also resulted in lower rates of survival *P*_time_ < 0.001; Fig. 1B), and flies scored as Smurfs were slightly, but significantly, more likely to die from a cold stress (*P*_Smurf_ = 0.024; Fig. 1B).

**Figure 1.**
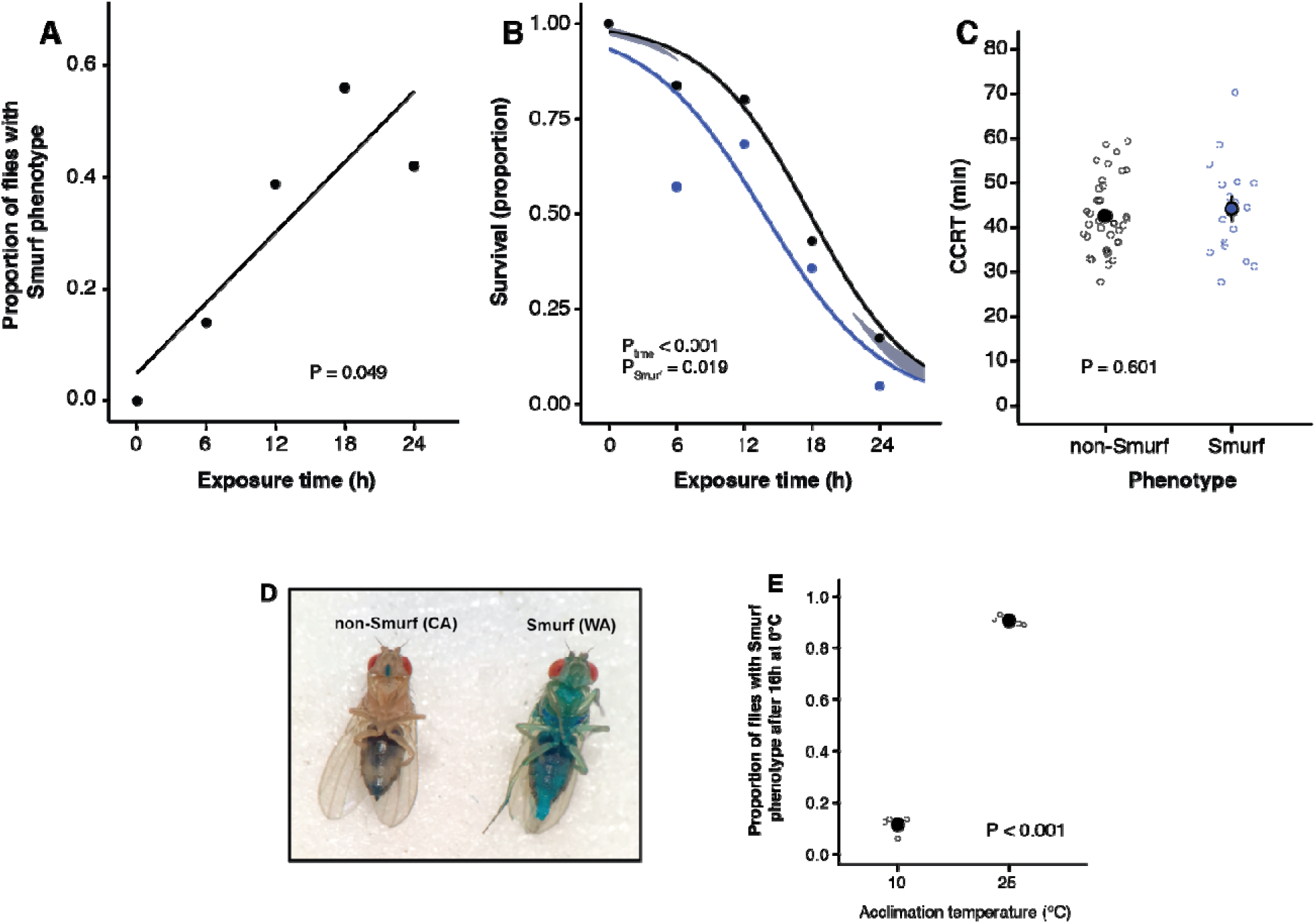
Cold stress causes the Smurf phenotype. A) Proportion of flies with a Smurf phenotype following exposure to 0°C for up to 24 h (n=49-50 flies per time period). B) Rates of survival of cold exposed flies that did, and did not adopt the Smurf phenotype during cold exposure. The time to cause 50% mortality at 0°C (Lt_50_) was slightly lower in Smurfs (13.8 ± 1.53 h) than in non-Smurfs (17.8 ± 1.18h). C) Chill coma recovery times (CCRT) of Smurf and Non-Smurf flies following 12 h at 0°C (n=56 flies in total; 17 Smurf, 39 non-Smurf). Open circles represent CCRT of individual flies and closed circles represent the mean ± sem. Recovery from chill coma was not related to the Smurf phenotype. D) Representative cold-(10°C) and warm-acclimated (25°C) flies following 16 h at 0°C. E) Proportions of cold- and warm-acclimated flies with a Smurf phenotype following 16 h at 0°C. Cold acclimation strongly reduces the probability of flies exhibiting a Smurf phenotype. Open circles represent proportions from n=4 replicates per acclimation temperature, each with 20-26 flies. Closed circles represent the mean ± sem. Error bars that are not visible are obscured by the symbols.

The chill coma recovery time (CCRT) of both Smurf and non-Smurfs was scored following 12 h at 0°C. Although the tendency to turn blue was related to low temperature survival, it was not significantly related to CCRT. The Smurf flies had an average recovery time of 44.2 ± 10.9 minutes, and the non-Smurf flies had an average recovery time of 42.6 ± 8.07 minutes (t_23_ = 0.530, *P* = 0.6010; Fig. 1C).

### Effects of acclimation on barrier disruption

To examine whether improvements in cold tolerance associated with cold acclimation may be related to the ability of the flies to avoid barrier disruption in the cold, we quantified the number of warm- and cold-acclimated flies with a Smurf phenotype following cold stress. Both warm- and cold-acclimated flies fed Bloomington food containing BB-FCF retained the dye in the gut lumen prior to any cold stress, but exposure to 0°C for 16h caused flies to lose barrier function and turn blue (Fig. 1C-E). The tendency to turn blue during cold stress, however, was significantly associated with the acclimation history of the flies (t_4_ = 39.2, P < 0.001); the cold exposure caused ∼ 90% of warm-acclimated flies to turn blue, only ∼ 10% of cold-acclimated flies turned blue after the cold stress (Fig. 1E).

### Smurf recovery assay

The Smurf recovery assay was used to test whether *Drosophila* with the Smurf phenotype could recover, and clear the BB-FCF from their hemolymph. If possible, we expected such recovery to change flies from a cold-induced Smurf to a non-Smurf phenotype within 30 h at benign temperatures. None of the control flies (i.e. those not given any cold exposure) became Smurf flies (n=25), and consequently, all of these control flies (n=25) were still non-Smurfs after the 12-hr recovery from the blue food, with 7 flies showing some residual blue dye inside the rectum. Of the total Smurf flies (n= 40), only 16 survived the cold exposure, however, of the 16 Smurf flies that survived, 14 resulted in non-Smurf phenotypes following the 30 h recovery period (Fig. 2). The remaining flies (n= 2) had some residual blue within their hemolymph and rectum.

**Figure 2.**
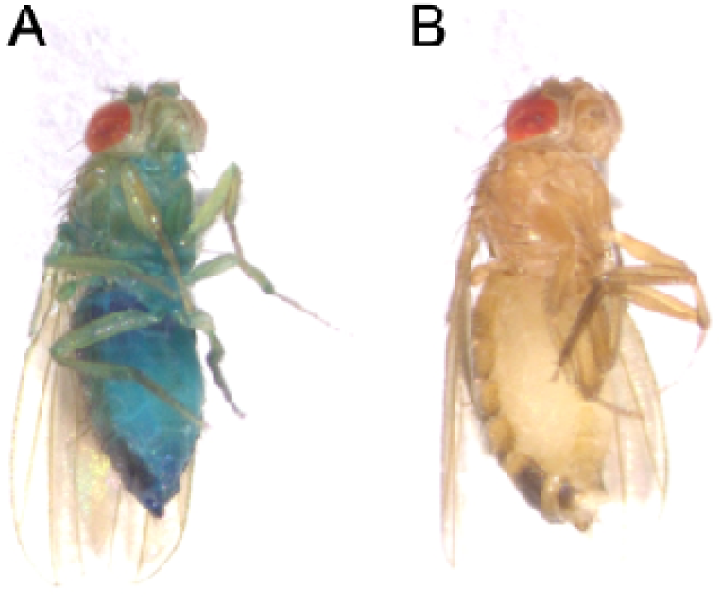
*D. melanogaster* can clear brilliant blue FCF from their hemolymph, reversing the Smurf phenotype. A) Smurf fly directly after removal from the cold stress. B) Smurf fly shown after recovery. Flies in 40 mL vials were submerged in an ice bath at 0°C for 12 hours to allow for the onset of the Smurf phenotype. The flies were then removed from the ice bath, and after inspection for the Smurf phenotype, they were put into an incubator at 25°C for 30 hours to allow them to clear the dye from the hemolymph.

### Midgut dye leak and transport assay

To test whether the midgut leaks BB-FCF *in situ*, we measured the appearance of the dye in a droplet surrounding the posterior midgut of flies. Notably, midguts of flies fed on the dye had measurable leak (3.32 ± 0.49 pmol mm^-1^ h^-1^ in the mucosal to serosal direction) even at room temperature (Fig. 3). Midguts also leaked the dye significantly faster at 0°C than at 23°C (F_14_ = 7.7, *P* = 0.015). To test whether control (room temperature) midguts could also transport the dye up a concentration gradient (despite concurrent leak), we added a known concentration of the dye to the bathing saline, and quantified disappearance of the dye from the saline. At 23°C, midguts had a net transport rate of the dye (in the serosal to mucosal direction) of 6.10 ± 0.66 pmol mm^-1^ h^-1^ (Fig. 3).

**Figure 3.**
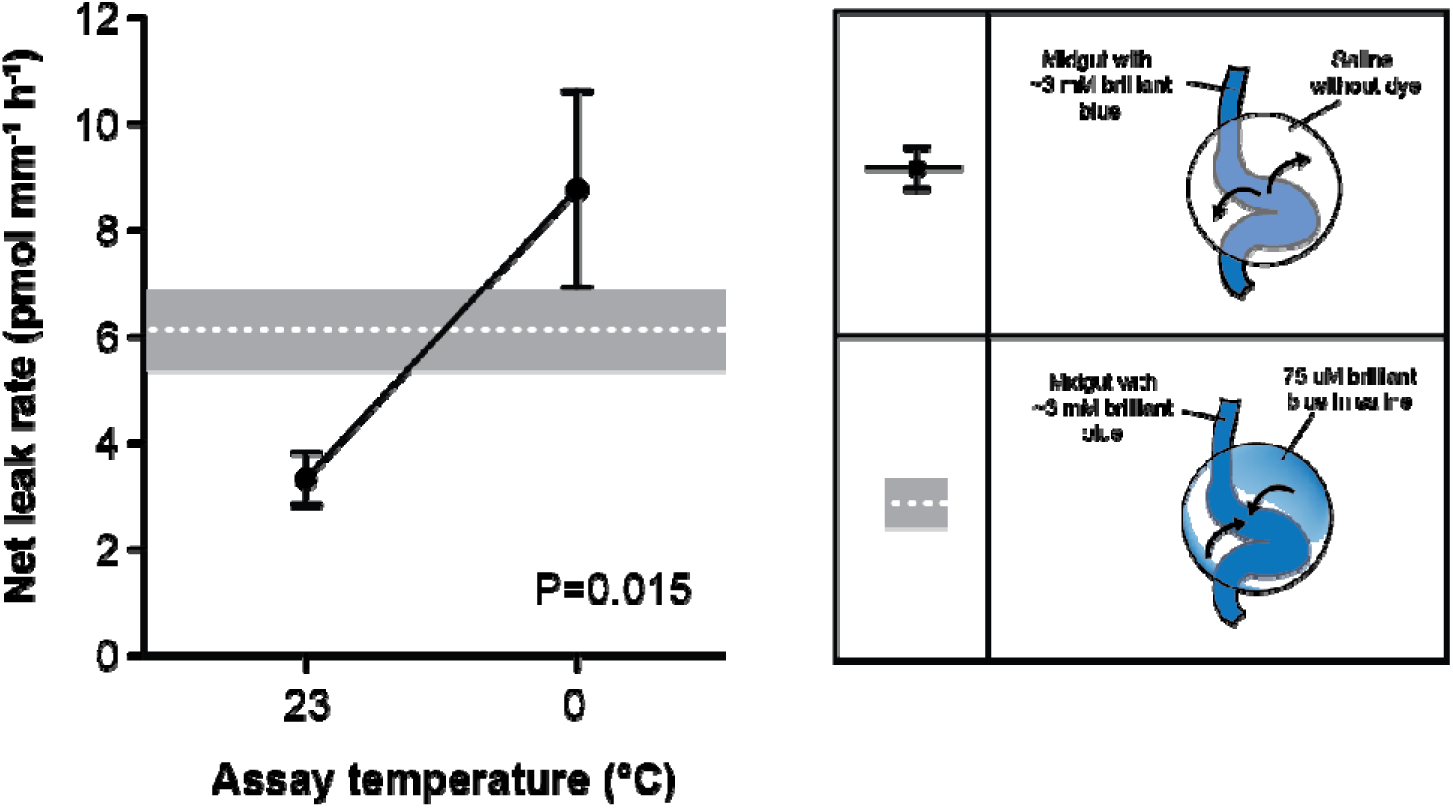
The midgut of *D. melanogaster* can transport brilliant blue FCF up a steep concentration gradient and cold causes net leak of the dye down its concentration gradient. Midguts dissected from adult female flies that had fed on 2.5% w/v brilliant blue FCF were submerged under oil and a droplet of saline containing no dye was placed on the posterior midgut and left for 3 h at either 23°C or 0°C. Net leak rates of the dye (solid circles; mean ± sem) were calculated from the appearance of the dye in the saline relative to the length of midgut tissue suspended in the droplet. These values are compared to the rate of net dye transport at 23°C, calculated from the rate of disappearance of the dye from a droplet containing a known concentration of brilliant blue FCF (75 µM; white dashed line and grey area; mean ± sem).

### Ramsay Assays

The Malpighian tubules of *D. melanogaster* were used to determine whether or not the tubules could actively transport BB-FCF. The Michaelis-Menten model closely fit the data and estimated a J_max_ value (highest possible rate of flux) of 232 fmol min^-1^, and a K_t_ value (point where rate of flux reached half of J_max_) of 760 µmol L^-1^ (Fig. 4). These results are also demonstrated qualitatively with example images of the Ramsay assays at each initial saline concentration in the supplementary material (Fig. S1).

**Figure 4.**
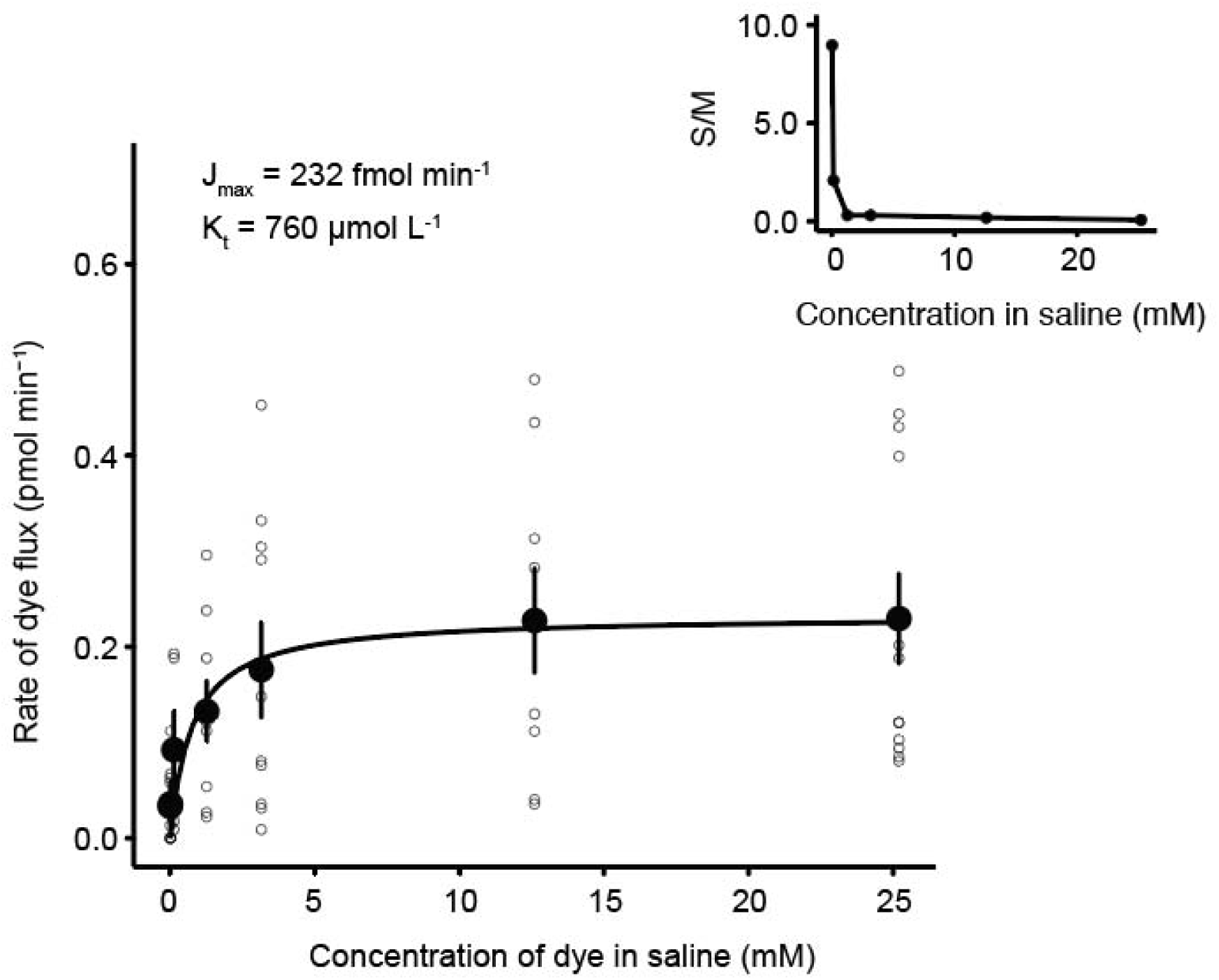
Rates of brilliant blue FCF transport through the Malpighian tubules of adult female *Drosophila melanogaster*. Modified Ramsay assays were used to characterize rates of transport of BB-FCF within 15 min of exposure to the dye. The black line denotes the fit of a Michaelis-Menten model. Solid symbols represent the mean ± sem and open circles represent values from individual tubules. Insert shows the ratio between the secreted BB-FCF concentration (S) and the initial bathing solution of the BB-FCF (M).

## Discussion

Here, we demonstrate for the first time that cold stress (like advanced age or traumatic injury) causes the Smurf phenotype in *Drosophila*, and that the tendency to turn blue is related to cold tolerance. We also show, however, that *Drosophila* appears to be able to transport BB-FCF, which complicates its use as a marker of barrier disruption.

Like other stresses, cold exposure caused flies to adopt a Smurf phenotype (turn blue), and longer cold stresses increased the proportion of Smurf flies (Fig. 1A). We noted an increase in abundance of Smurf phenotypes as the duration of cold exposure increased and as survival rates declined, but there was also a drastic decrease in survival as well (Fig. 1B). With these findings alone, one could presume that barrier failure was being well characterized by this blue dye, especially considering that barrier dysfunction occurs in the cold when measured using dextran (MacMillan et al., 2017), and that other large polar molecules like glucose can leak into the hemolymph under conditions of stress (Katzenberger et al., 2015).

Insects, including *Drosophila* have an acute ability to recover from cold exposure (Hoback and Stanley, 2001). Upon chilling, *Drosophila* will enter a comatose state (chill coma), wherein their neural and muscular systems are silenced (Rodgers et al., 2010). After they return to a normal homeostatic temperature, the flies can revert from the comatose state back to a functional state (David et al., 1998). The time required to return from chill coma is a common measure of cold tolerance, and is related to the degree to which ion and water balance have been disturbed, both in the nervous system and the hemolymph (Andersen and Overgaard, 2019). We performed a CCRT experiment using the Smurf flies and the non-Smurf flies, and found no difference in CCRT between the two phenotypic groups (Fig. 1C). This suggests that the severity of ion balance disruption is not related to the tendency of BB-FCF to leak across the epithelial barrier. Therefore, the cold stress appeared to make the epithelial barriers leaky, and this leak was associated with low temperature, death and injury, but not with chill coma recovery.

We demonstrated that under normal conditions, there was no evidence of BB-FCF leaking/being transported into the hemolymph, however after a stress is applied, there is strong evidence of net leak of BB-FCF. Thermal acclimation can drastically alter cold tolerance phenotypes in adult *Drosophila* (Ransberry et al., 2011; Colinet and Hoffmann, 2012). To examine whether or not the tendency for flies to turn blue was related to thermal plasticity, we then investigated whether or not acclimation would similarly impact the tendency of flies to turn blue during a cold stress. Using this approach, we found a striking reduction in the tendency for cold-acclimated flies to accumulate the blue dye in their hemolymph, relative to warm-acclimated flies (Fig. 1D-E). We suggest two possible explanations for a reduction in Smurf phenotypes given a cold acclimation; 1) cold acclimation prevents epithelial barrier failure (MacMillan et al., 2017), or 2) cold acclimation improves the ability of transporters to clear the dye from the hemolymph in the cold. Given these two possibilities we opted to investigate whether or not flies were capable of clearing BB-FCF from their hemolymph.

Our Smurf recovery assay demonstrated the ability of the flies to revert their phenotype from Smurf to non-Smurf, which suggests that the flies can either transport and excrete BB-FCF out of their hemolymph, or they are able to effectively metabolize it. In order to clarify whether flies are capable of transporting BB-FCF, we tested for the capacity for transport of this dye through the midgut and/or the Malpighian tubule epithelia using *in situ* assays. We took this approach because BB-FCF dye is an organic anion which is similar in structure to other sulphonates (e.g. amaranth, indigo carmine) that are already known to be transported by the Malpighian tubules and midgut of *D. melanogaster* at relatively high rates (Linton & O’Donnell, 2000; O’Donnell et al., 2003). We found that the *Drosophila* midgut is capable of effectively transporting this dye against a concentration gradient (Fig. 3). This is similar to other large organics, which have also been found to be transported across the midgut epithelium (Rheault & O’Donnell, 2004, Bijelic et al., 2005; Chahine & O’Donnell, 2009). The Malpighian tubules function in ion and water regulation but are also critical to the clearance of wastes and toxic dietary compounds from the hemocoel (Dow, 2009; Denholm, 2013). We investigated the ability of the Malpighian tubules to transport BB-FCF via modified Ramsay assays (Rheault and O’Donnell, 2004). The rate of flux of BB-FCF into the Malpighian tubules was seen to increase along with the bathing concentration at low concentrations (Fig. 4), but plateaued at high concentrations. A Michaelis-Menten model was fit to the relationship between the concentration of the dye in the bathing saline and the rate of dye flux into the primary urine (Fig. 4). We found that the J_max_ of BB-FCF is approximately 232 fmol min^-1^, which is comparable to other organic compounds such as methotrexate and Texas red which have J_max_ values of 118.4 and 1028 fmol min^-1^ respectively (Leader and O’Donnell, 2005; Chahine and O’Donnell, 2009). Tetraethylammonium (TEA) is similarly sized but a cation (and is thus mobilized by different transporters) but has a much higher J_max_ of 1.52 pmol min^-1^ tubule^-1^ (Rheault and O’Donnell, 2004). The K_t_ of BB-FCF (760 µM) was higher than those of methotrexate and TEA in the Malpighian tubules of *D. melanogaster*, as the K_t_ values for these two molecules were 12.2 µM and 220 µM respectively (Rheault and O’Donnell, 2004; Chahine and O’Donnell, 2009). Overall, our Ramsay assay results indicate that transport of BB-FCF is saturable, like several other similarly-sized organic compounds.

There are two known types of organic anion transporters in the *Drosophila* Malpighian tubules; MRP1 and MRP2 (Multidrug resistant proteins) that transport either type I or type II organic anions (OAs) (Leader & O’Donnell, 2005; Chahine et al., 2012). Type I OAs are generally smaller molecules and are transported by Na^+^-dependent processes (Bresler et al., 1990; Linton & O’Donnell, 2000). Type II OAs are typically larger molecules and are transported Na^+^- independently (Leader and O’Donnell, 2005; Chahine and O’Donnell, 2009). Texas red, which is an organic anion of a similar size to BB-FCF, is transported by MRP2 (Leader and O’Donnell, 2005). Based on similarity to Texas red, we suggest that BB-FCF is actively transported in a Na^+^-independent manner by MRP2. In support of this hypothesis, BB-FCF has recently been shown to inhibit MRP2 transport in human intestinal membrane vesicles in substrate competition experiments, and MRP2 is capable of transporting BB-FCF into vesicles (Sjöstedt et al., 2017). Future studies could verify whether MRP2 is responsible for the transport of BB-FCF into the Malpighian tubules, such as transgenic knockdown of MRP2 or substrate competition experiments.

A good marker of paracellular permeability requires a few important qualities; it should 1) have a distinct colorimetric or radioactive attribute (allowing the marker to be quantified), 2) be non-lethal to the organism, and 3) not be able to be transported by the tissues and organs of the organism under study. We argue that brilliant blue FCF, while convenient in studies of barrier disruption for the first two requisite attributes, violates the third requirement. In most of the present studies using BB-FCF to characterize barrier dysfunction, the non-Smurfs are presumed to maintain all of the brilliant blue dye in the gut unless gut failure occurs, but we show here that this may not be the case. Based on our findings, the midgut appears instead to leak the dye out of the gut into the hemolymph, and the gut and Malpighian tubules secrete the dye back into the gut in a constant cycle. Thus, even if a fly fed the dye is not a Smurf, there is still some brilliant blue in the hemolymph at all times. The dye is simply being actively transported back into the gut at high enough rates so as to keep the concentration of BB-FCF in the hemolymph low enough to evade visual detection (i.e. all the flies are Smurfs, but some are more Smurfy than others). Based on this model, any break in the cycle (e.g. via an inhibition of transport, paracellular barrier failure, or physical damage to the gut) could induce the Smurf phenotype, and observation of the organismal phenotype is not indicative of any one form of failure. In the case of cold exposure, low temperatures supress active transport rates by a factor of 2-3 for every 10°C drop in temperature (the Q_10_ effect), and cold causes paracellular barrier disruption (MacMillan et al., 2017), so at least two factors likely contribute to the cold-induced Smurf phenotype described here.

We examined the practical use of brilliant blue dye (BB-FCF), and whether or not *Drosophila* could transport it via their midgut and/or Malpighian tubules. We concluded that both the midgut and Malpighian tubules have the ability to transport the dye. These results together suggest that BB-FCF is not an effective tool for measuring paracellular barrier dysfunction, as *Drosophila* can and do effectively transport BB-FCF across their intestinal epithelial barriers, possibly via MRP2. Importantly, these findings should not be interpreted as undermining the importance of the Smurf assay or what has been learned from it. Instead they should be seen as allowing for new hypotheses on the precise mechanisms that cause a fly to turn blue during stress or at an advanced age. In addition, we argue that the insect community should actively test other methods of assaying paracellular barrier disruption, and be careful to avoid the assumption that markers used in vertebrates (e.g. dextran, inulin, or polyethylene glycol) are inherently good markers for insects.

## Supporting information

Data archive

## Data Accessibility

All data is provided as a supplementary file for review and the same file will be uploaded to a data repository (e.g. Dryad) should the manuscript be accepted for publication.

## Author Contributions

All authors contributed to the conception and design of the study, D.L. H.P., and H.M. conducted the experiments. D.L. and H.M. analyzed the data, D.L. drafted the manuscript, and all authors edited the manuscript.

---

**Figure S1.**
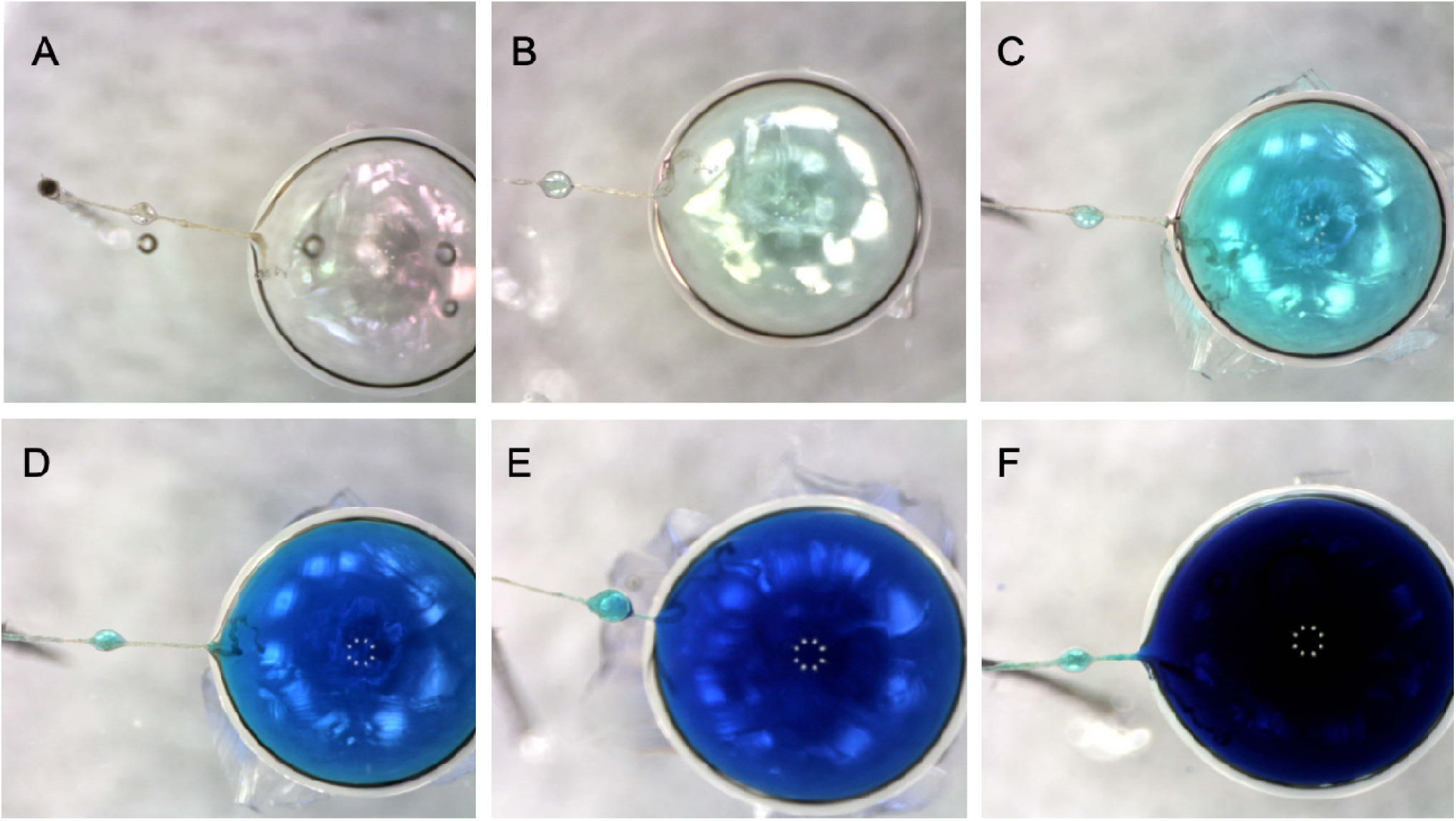
The *Drosophila* Malpighian tubules can transport brilliant blue FCF. Modified Ramsay assays were performed to examine the effects of increasing BB-FCF concentrations in the bathing saline on the concentration of the resultant primary urine. A = Control (0 µM), B = 0.0126 µM, C = 0.126 µM, D = 1.26 µM, E = 3.15 µM, and F = 12.6 µM.

